# Predictive processing in the cortical network of biological motion perception: An fMRI-DCM study

**DOI:** 10.1101/2025.03.05.641401

**Authors:** Murat Batu Tunca, Burcu A. Urgen

## Abstract

Biological motion perception (BMP) is fundamental for interpreting and predicting human actions, playing a crucial role in social cognition. While traditionally viewed as a bottom-up process, recent evidence suggests that top-down mechanisms, particularly expectations, influence BMP. This fMRI study investigates how prior expectations shape BMP by employing a predictive cueing paradigm. Participants discriminated between biological and scrambled motion while presented with congruent, incongruent, or uninformative cues. Behavioral results revealed that correct expectations facilitated performance, while incorrect expectations impaired it. Dynamic Causal Modeling (DCM) analysis demonstrated bidirectional connectivity within the action observation network (AON), particularly between the posterior superior temporal sulcus (pSTS), posterior parietal cortex (PPC), and inferior frontal gyrus (IFG) under all expectation conditions. Additionally, Multivariate Pattern Analysis (MVPA) confirmed the involvement of these regions in BMP. These findings challenge traditional bottom-up models by highlighting the role of predictive processes in action perception and provide evidence for expectation-dependent modulation of AON connectivity.

## 1. Introduction

Biological motion perception (BMP) is a vital cognitive skill for animal survival. In humans, it serves as the foundation of successful social interactions in everyday life. Previous studies in the field have shown that BMP is distinct from general motion perception (Chouchourelou et al., 2013). It relies on a specialized neural network known as the action observation network (AON) consisting of the posterior superior temporal sulcus (pSTS), posterior parietal cortex (PPC), and inferior frontal gyrus (IFG) (Jastorff & Orban, 2009; Peuskens et al., 2005; Lesourd et al., 2022; Orban, 2018; Grossman & Blake, 2013; Blake & Shiffrar, 2007; Calvo-Merino et al., 2005; Cross et al., 2006; Grezes & Decety, 2001; Saygin et al., 2004; van Kemenade et al., 2012). Albeit not a part of the AON, some studies have reported selectivity for biological motion in the fusiform gyrus as well (Vaina et al., 2001). Most commonly, biological motion studies are conducted by using point-light displays (PLDs). These controlled stimuli are enough to activate the AON (Johansson, 1973; Blake & Shiffrar, 2007; Troje, 2002). Similarly, a standardized control stimulus in the literature is to use the scrambled versions of these PLDs (Bonda et al., 1996).

One common finding among multiple biological motion studies was that participants were highly fast and accurate in the perception of biological motion, even in tasks with significant noise (Blake & Shiffrar, 2007; Freitag et al., 2008). The ease in its processing has led researchers to believe that biological motion perception is an automatic process modulated solely by bottom-up information. Supporting this idea, the researchers demonstrated that participants were able to determine the motion direction when only the feet were visible to them (Chang & Troje, 2009; Saunders et al., 2009). This finding was explained as the motion of the feet was triggering the necessary innate cognitive mechanisms for biological motion perception, recognition, and interpretation (Chang & Troje, 2009; Saunders et al., 2009).

The importance of bottom-up processes in biological motion perception also found its place in theoretical models. The original model presented by Johansson (1973) proposes that biological motion perception is a hierarchical mechanism that utilizes solely feedforward information. It argues that minimal and simple motion cues are integrated into the perception of biological motion (Johansson, 1973). Similarly, Giese and Poggio’s model (2003) emphasizes that low-level motion cues are integrated into more complex information, which is then compared to specific motion templates. These templates, encoded in neurons specifically tuned for motion perception, allow the correct perception and interpretation of the biological motion.

As biological motion literature progressed, researchers started to challenge the view that biological motion perception is solely a bottom-up process. In everyday life, one’s perception is constantly affected by top-down processes, most notably attention and expectation. To investigate the role of attention, studies used Lavie’s (1995) attentional load theory as a basis. The theory states that attention is a limited resource and multiple stimuli continuously compete for one’s attention. Aligning with Lavie’s theory, studies have shown that attending to a biological motion stimulus significantly facilitates its processing. On the other hand, when participants are unable to attend to the biological motion because their attention is directed toward a high-load task, their perception of biological motion will be noticeably impaired or even prevented (Thompson et al., 2005; Safford et al., 2010; Gilaie-Dotan et al., 2013; Hars et al., 2011; Parasuraman et al., 2009; Tunca et al., 2023; Nizamoğlu & Urgen, 2023; Battelli et al., 2003; Cavanagh et al., 2001; Thornton et al., 2002; Pavlova et al., 2006; Neri et al., 1998).

While the effect of attention on biological motion perception has been investigated, the effect of expectations remains largely unexplored. Nevertheless, expectation literature proposes significant evidence to demonstrate how prior knowledge shapes perception. The review by de Lange and colleagues (2018) provides numerous examples of how prior information is combined with sensory input to form, alter, or impair the perception of stimuli. Behavioral and neuroimaging studies further demonstrate that accurate prior information enhances processing efficiency, with the prefrontal cortex and medial temporal lobe implicated in generating predictions (Kok et al., 2014; Kok et al., 2012; Kok et al., 2013; Miller & Cohen, 2001; Schacter et al., 2007). When these predictions are violated, a prediction error signal emerges, involving the ventral tegmental area (VTA), anterior cingulate cortex (ACC), striatum, and prefrontal cortex (Miller & Cohen, 2001; Schultz, 1998; Behrens et al., 2007; Schultz, 2015; Jessup et al., 2010).

These experiments provide strong evidence that expectations shape our perception. However, it should be acknowledged that expectation studies have mostly utilized low-level and simple stimuli. For visual expectation studies, the stimuli presented are often Gabor filters or moving dots. While the utilization of such elementary stimuli helped isolate neural mechanisms of expectation, they lacked considerable external validity. Ultimately, the effect of expectation on more complex and social stimuli remains mostly unexplored. The current study aims to fill this gap by investigating the effect of expectation on a highly complex and socially meaningful stimulus, the biological motion.

One of the few studies that aimed to fill this gap has revealed that the action observation network’s activity depends on prediction violations, suggesting the generation of prediction errors (Saygin et al., 2012). This finding aligned with the more modernized view of AON as a predictive coding system that consists of both feedforward and feedback connections, challenging the bottom-up models of biological motion perception (Friston, 2010). The view found further support in anatomical brain imaging studies that revealed reciprocal connections between PPC and pSTS, and between PPC and IFG (Luppino et al., 1999; Seltzer & Pandya, 1994; Rushworth et al., 2006; Tgelstrom & Graziano, 2017). Urgen & Saygin (2020) extended this finding by investigating the effective connectivity between AON nodes. Their study has demonstrated that when a mismatch occurs between the look and the movement of an agent, prediction errors are created in PPC and are sent to the premotor cortex (Urgen & Saygin, 2020). Similarly, Elmas and his colleagues’ (2024) behavioral study has found that biological motion perception is hindered when a mismatch between expected and observed actions is present (Elmas et al., 2024).

Building on these insights, the current study aims to investigate a common real-life scenario: expecting a specific action from someone but observing a different one instead. Specifically, we examine the neural mechanisms underlying the mismatch caused by unexpected actions. To achieve this, we employ Dynamic Causal Modeling (DCM), an effective connectivity technique that enables the examination of the direction and strength of connections between AON regions (Friston et al., 2003, Penny et al., 2004). Understanding these connections is crucial, as it provides empirical support for the AON as a predictive coding system (Friston, 2010). Furthermore, identifying feedback connections within the network would challenge bottom-up models of BMP and contribute to the development of a novel framework. Additionally, connectivity analyses will clarify whether interactions within the AON are excitatory or inhibitory and how they depend on prediction processes.

By integrating research on BMP and predictive coding, this study aims to elucidate the influence of expectations on action perception. Participants are presented with a paradigm in which biological motion is preceded by a cue indicating the expected action type. In line with prior research, we anticipate behavioral and neural differences between conditions where predictions are upheld versus violated. Through this approach, we seek to advance our understanding of how expectation modulates BMP and its underlying neural mechanisms.

## 2. Methods

### 2.1 Participants

fMRI data of 29 participants was collected (Range = 18-30, Mean = 22.6, SD = 3.0, 17 female, 12 male). Before the experiment, the participants were fully informed about the procedure and gave their informed consent approved by the Human Research Ethics Committee of Bilkent University. After the experiment, the participants were compensated with either money (50 TL) or course credit, depending on their choice.

### 2.2 Stimuli

Two types of stimuli were used in the experiments: Cue stimuli and point-light displays. The cue stimulus was always a static image and had three variations: A walking man, a kicking man, or a question mark. Similarly, the point-light displays had four variations: The motion of a kicking man, the scrambled version of the kicking man, the motion of a walking man, and the scrambled version of the walking man. Each point-light display consisted of 13 dots. The starting points of each dot from kicking man and walking man motions were randomized to generate their scrambled versions. (Figure 2.1). Although other possible actions were considered, the study decided to use walking and kicking stimuli for two main reasons. First, the importance of foot information for biological motion perception was shown by previous studies (Freitag et al., 2008; Chang & Troje, 2009). Second, research has shown that the processing of communicative actions differs from the processing of non-communicative actions (Manera et al., 2011; Urgen & Orban, 2021). Thus, since the current study is only interested in the processing of motion information independent of communicative intent, non-communicative actions were selected. Ultimately, walking and kicking actions, two non-communicative actions with clear but distinct foot movements, were found to be ideal for the current study. The cue stimuli (3.5 x 3.5 degrees in size) were presented at the center of the screen. On the other hand, the point light displays (5.2 x 8.3 degrees in size) were presented at 6 degrees to the left or right of the center.

**Figure 2.1.**
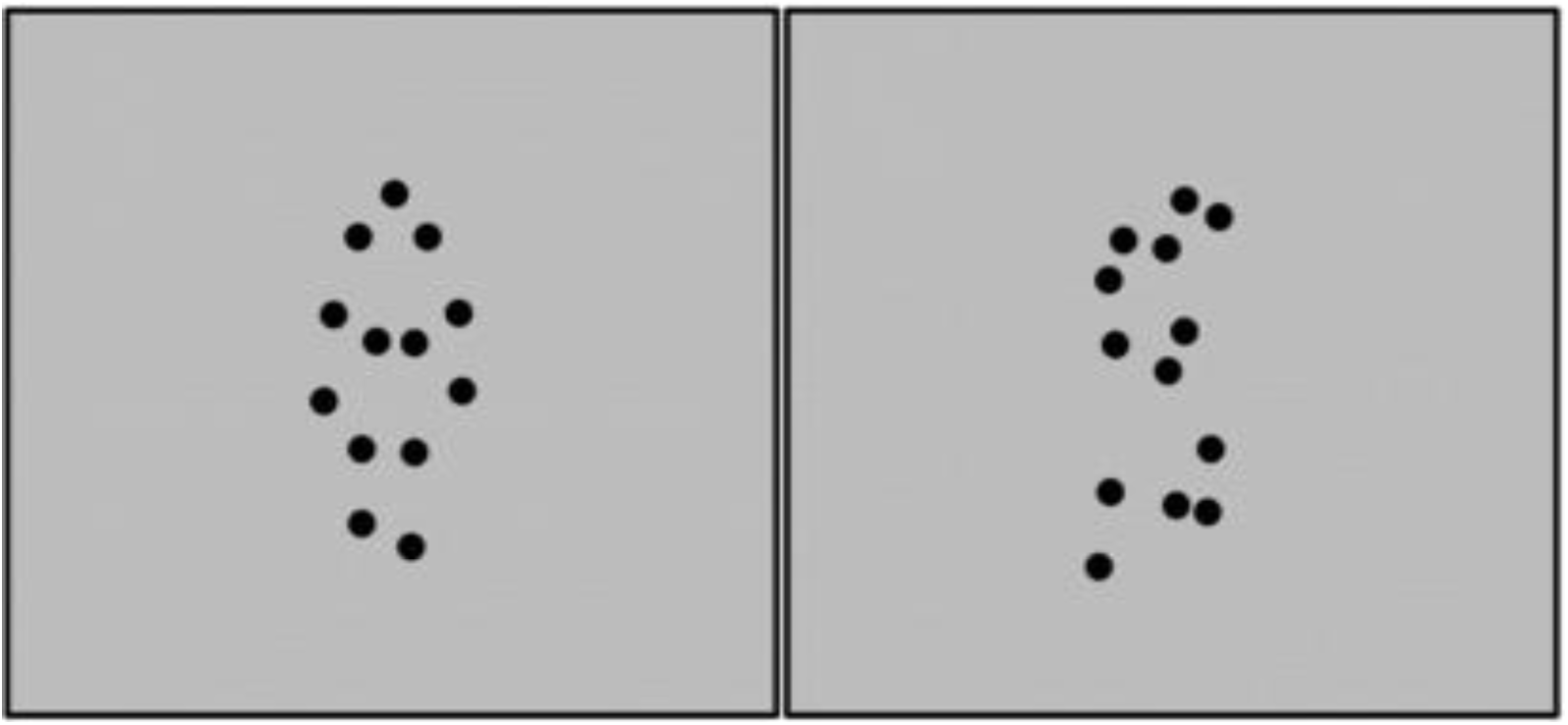
Point-light Displays. *Note.* Figure at the left: Biological motion representing a walking man. Figure at the right: Its scrambled version.

### 2.3 Experimental Design and Procedure

The experiment was coded in MATLAB with the help of PsychToolBox and Biomotion Toolbox (Brainard, 1997; Pelli, 1997; Boxtel & Lu, 2013). Each run of the experiment consisted of four conditions: Congruent, incongruent, no cue (neutral), and blank. The steps of a congruent trial were as follows (Figure 2.2.A): The trial would start with the presentation of the walking man or kicking man cue for 2 seconds. This was followed by an interstimulus interval of 0.5 to 1.5 seconds. Then, two point-light displays (one on the left and one on the right) were presented for 1.7 seconds. The action performed by the displays was consistent with the cue, however, one of the displays was the biological motion whereas the other one was its scrambled version. This presentation was followed by the response period for 2 seconds during which the participants were asked to indicate on which side of the screen (left or right) the biological motion (not the scrambled motion) was presented. The participants reported their responses using the left and right buttons of an MRI-compatible button box. Then, the participants were given feedback about the accuracy of their responses for 1 second. After the feedback period, another interstimulus interval for 3 to 4 seconds would mark the end of the trial. In total, each trial lasted between 10.2 to 12.2 seconds. Additionally, throughout the experiment, the participants were informed to focus on the fixation point presented at the center of the screen as a black plus sign. After the motion stimuli, the fixation turned white for 2 seconds to indicate to the participants that they were in the response period.

**Figure 2.2.**
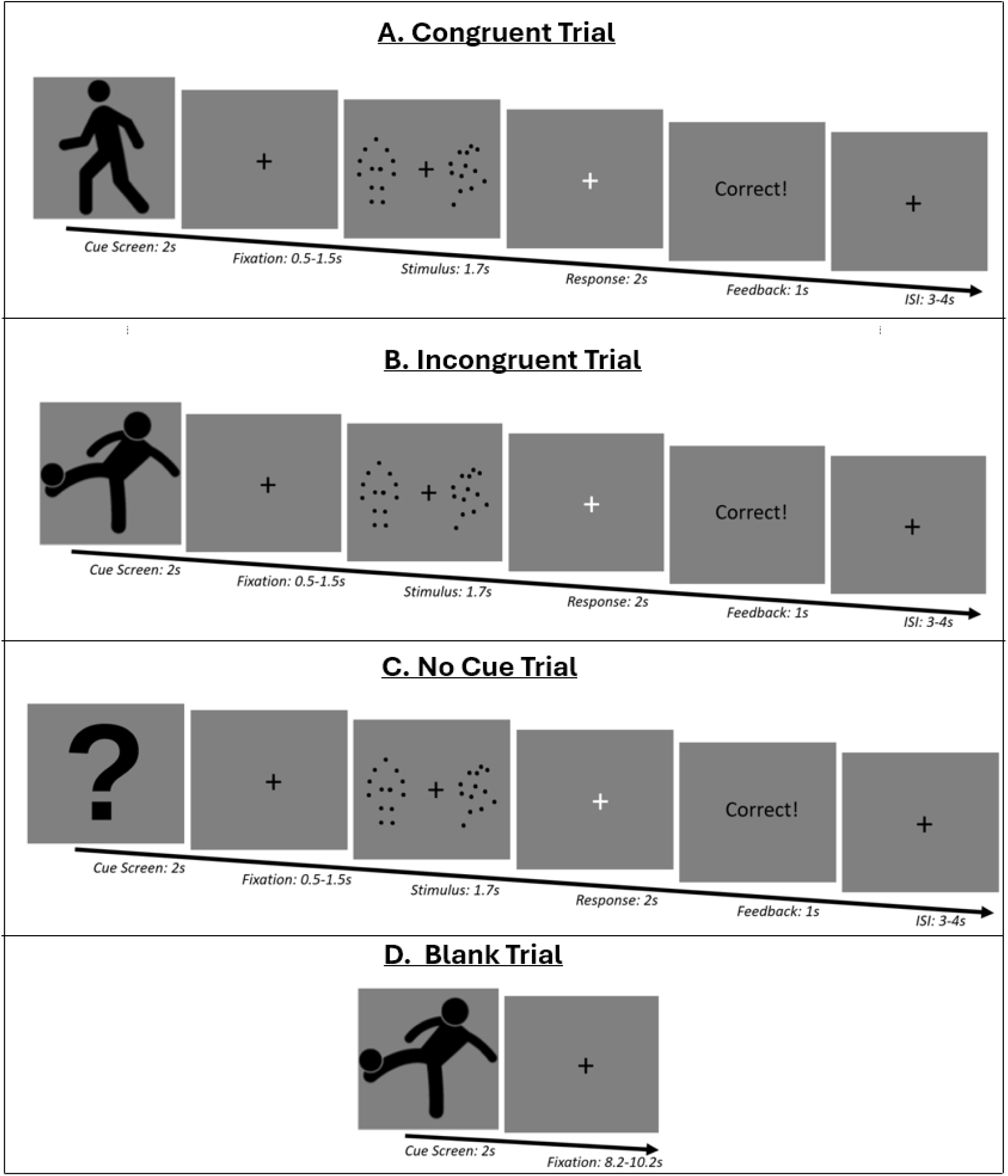
An example of each trial type. *Note.* **A.** A congruent trial with walking cue and walking PLD. **B.** An incongruent trial with kicking cue and walking PLD. **C.** A no cue trial with walking PLD **D.** A blank trial with kicking cue.

Other conditions of the experiment would follow the same steps as the congruent condition but with minor differences. During the incongruent condition, the point light displays presented to the participant were inconsistent with the presented cue (Figure 2.2.B). On the other hand, during the no cue condition, the cue was an image of a question mark so that it would give no information about the following displays (Figure 2.2.C). During the blank trials, a cue was presented to the participants but no display followed it. Instead, the participants saw the fixation point for 8.2 to 10.2 seconds (Figure 2.2.D). The participants were informed that no response was expected from them during blank trials.

The experiment was designed to have a 75% congruence rate between the cue and the displays. To achieve this, each run included 12 congruent trials and 4 incongruent trials. The control conditions, which were no cue and blank conditions, were presented in the same amount as the incongruent trials. Within each condition, each possible cue-stimulus pairings were presented an equal number of times (i.e. walk-walk and kick-kick pairings were each presented 6 times, and kick-walk, walk-kick, ?-kick, ?-walk, walk-blank and kick-blank were each presented twice). Also, each run started and ended with a fixation period of 10 seconds. Ultimately, each run included 24 trials and lasted for 5 minutes. The experiment spanned 10 runs and the participants were free to rest their eyes as long as they wanted between runs.

In order to prevent ceiling performances, randomly moving noise dots were added to the displays. Before the experiment, the participants were asked to complete a short behavioral practice period to determine their optimal noise level and to make sure that they understood the task. After giving detailed instructions about the task, the biological motions and scrambled motions were introduced to the participant. Then the first practice experiment started, which was an 80-trial version of the original experiment. This version was slightly different from the original experiment with shorter interstimulus intervals (ranging from 0.8 to 1.2 seconds) and without the blank condition. During this practice session, the Quick Estimation by Sequential Testing (QUEST) method was used to adaptively calculate the ideal amount of noise level. The optimal noise level obtained at the end of this session would be unique to each participant and be utilized throughout the fMRI experiment.

Once the participants were done with this practice, they moved on to the second practice session. This version of the experiment consisted of 14 trials and exactly replicated the original experiment in terms of trial durations, noise level, and blank conditions. Before starting each practice session and the fMRI experiment, the participants were instructed that their priority should be to respond as accurately and quickly as possible, with an emphasis on accuracy.

### 2.4 fMRI Data Collection

A 3T Siemens TimTrio MRI scanner and 32-channel head coil in the National Magnetic Resonance Research Center (UMRAM, affiliated with Bilkent University) were used for the scanning periods. Foam sponges were placed under the heads, on the sides of necks, and under the legs of the participants to prevent them from moving during the experiment. The participants were able to see the MRI-compatible LCD screen (TELEMED, 60Hz refresh rate, 800x600 pixels, 32 inches) through a mirror placed on top of the head coil. The screen was positioned 168 cm away from the participants. Before the experiment began, high-resolution T1-weighted anatomical scans of the whole brain were taken (TE = 2.92ms, TR = 2.6s, flip angle = 12°, acceleration factor = 2, 176 sagittal slices with a resolution of 1mm x 1mm x 1mm). Functional scans were acquired using EPI sequences during the experiment (TE = 30ms, TR = 2s, flip angle = 90°, 240 field of view, 96x96 matrix, 49 horizontal slices with a thickness of 2.5mm). Each scanning session started with 3 dummy scans to ensure the scanner reached a stable level.

### 2.5 fMRI Data Analysis

#### 2.5.1 Preprocessing

Preprocessing was conducted using fMRIPrep 20.1.1, based on Nipype 1.5.0 (Esteban et al., 2019; Gorgolewski et al., 2011). Preprocessing steps were motion correction, slice-timing correction, registration, and normalization. Motion correction was performed using mcflirt by estimating head movements in 3 dimensions and 2 directions (Jenkinson et al., 2002). Slice-time correction was performed using 3dTshift from AFNI 20160207 (Cox & Hyde, 1997). Registration was performed using bbregister, with 6 degrees of freedom (Greve & Fischl, 2009). ICBM 152 Nonlinear Asymmetrical template version 2009c was selected for spatial normalization (Fonov et al., 2009).

#### 2.5.2 Dynamic Causal Modeling (DCM) Analysis

Dynamic Causal Modeling analysis was conducted to explore the strength and direction of connections between regions of interest (Friston et al., 2003; Penny et al., 2004). The analysis included two stages: Model specification and model selection. In the model specification stage, models were created with various connections depending on the previous literature and current hypotheses. For each model, endogenous and modulatory connections between regions of interest were estimated. This stage was followed by the model selection stage, which included Bayesian Model Selection procedure to calculate each model’s probability to explain the current data. The most probable model was declared as the winning model.

##### 2.5.2.1 Definition of Regions of Interest

Before defining ROIs, the data had to be optimized for DCM analysis. This was done by excluding certain runs or participants from the analysis. 8 participants’ data failed to have 10 runs as 4 participants asked to end the experiment prematurely after 8 runs and 5 participants (one of which also asked to end the experiment early) had 1 or 2 of their runs excluded from fMRI analysis because of head motion. Since DCM analysis required the same number of runs among participants, it was decided that only the first 8 runs of each participant would be included in the DCM analysis. ROIs were defined by using the first 8 runs of each participant. They were defined in 23 participants, excluding 6 participants. These participants were excluded from the analysis as one participant’s data contained only 7 runs, two participants’ data were invalid because of head motion and three participants’ data were invalid because of a technical problem with the MRI scanner.

A general linear model was conducted to define regions of interest. 11 regressors were used for this GLM. They were, in order: Congruent condition, incongruent condition, no cue condition, blank condition, and rest. The last 6 regressors were head movements of the participant in 3 dimensions and 2 directions. For each participant, a contrast was created by subtracting the blank condition from the combination of congruent, incongruent, and no cue conditions. This contrast of motion conditions with the non-motion condition was decided to be used for defining ROIs. Since the aim of DCM analysis was to investigate the connections within the action observation network, nodes of AON (pSTS, PPC and IFG) were pre-determined as ROIs. Therefore, the activation of created contrast was investigated in these nodes. Indeed, the second-level GLM analysis showed activation at the inferior frontal gyrus (IFG), posterior superior temporal sulcus (pSTS), and posterior parietal cortex (PPC). Although the activation was bilateral, activation at the right nodes was stronger compared to the left nodes. Therefore, the following brain regions were defined as ROIs: Right IFG (*x* = 54*, y* = 12*, z* = 27), right pSTS (*x* = 48*, y* = *−*52*, z* = 4), and right PPC (*x* = 27*, y* = *−*55*, z* = 54)(*p <* 0.01, not corrected*, k* = 0 for all). Then, a 5mm radius sphere was created at each ROI for each participant. The local maxima within this sphere served as the center point for a second sphere of 4mm radius, specific to each participant. Principal eigenvariate of all voxels that survived *p <* 0.01 threshold within that second sphere was used to extract the time series for each run. Two more participants had to be excluded from DCM analysis as their data failed to show activation in the selected ROIs.

##### 2.5.2.2 Model Specifications

In the model specification stage, to explain the current data, 64 models were created. These models were distinct from each other in terms of how connections between pSTS, PPC, and IFG were modulated by conditions of the study (congruent, incongruent, and no cue). In line with previous literature, all models assumed that information first enters the pSTS and a feedforward flow is present from pSTS to PPC and from PPC to IFG. On the other hand, the feedback connections were different among models. The general model, which included every possible connection of interest in the study, proposed that all 6 possible feedback connections were present alongside the feedforward connections. These feedback connections were: The feedback connection from PPC to pSTS that was modulated by congruent condition, the feedback connection from PPC to pSTS that was modulated by incongruent condition, the feedback connection from PPC to pSTS that was modulated by no cue, the feedback connection from IFG to PPC that was modulated by congruent condition, the feedback connection from IFG to PPC that was modulated by incongruent condition, and the feedback connection from IFG to PPC that was modulated by no cue condition. Every other model was then obtained by removing at least one feedback connection from the general model, resulting in 64 possible models. Then, the models were grouped into 8 families. Connections from PPC to pSTS were different for models among different families whereas connections from IFG to PPC were different for models within the same family (Table 2.1).

**Table 2.1.**
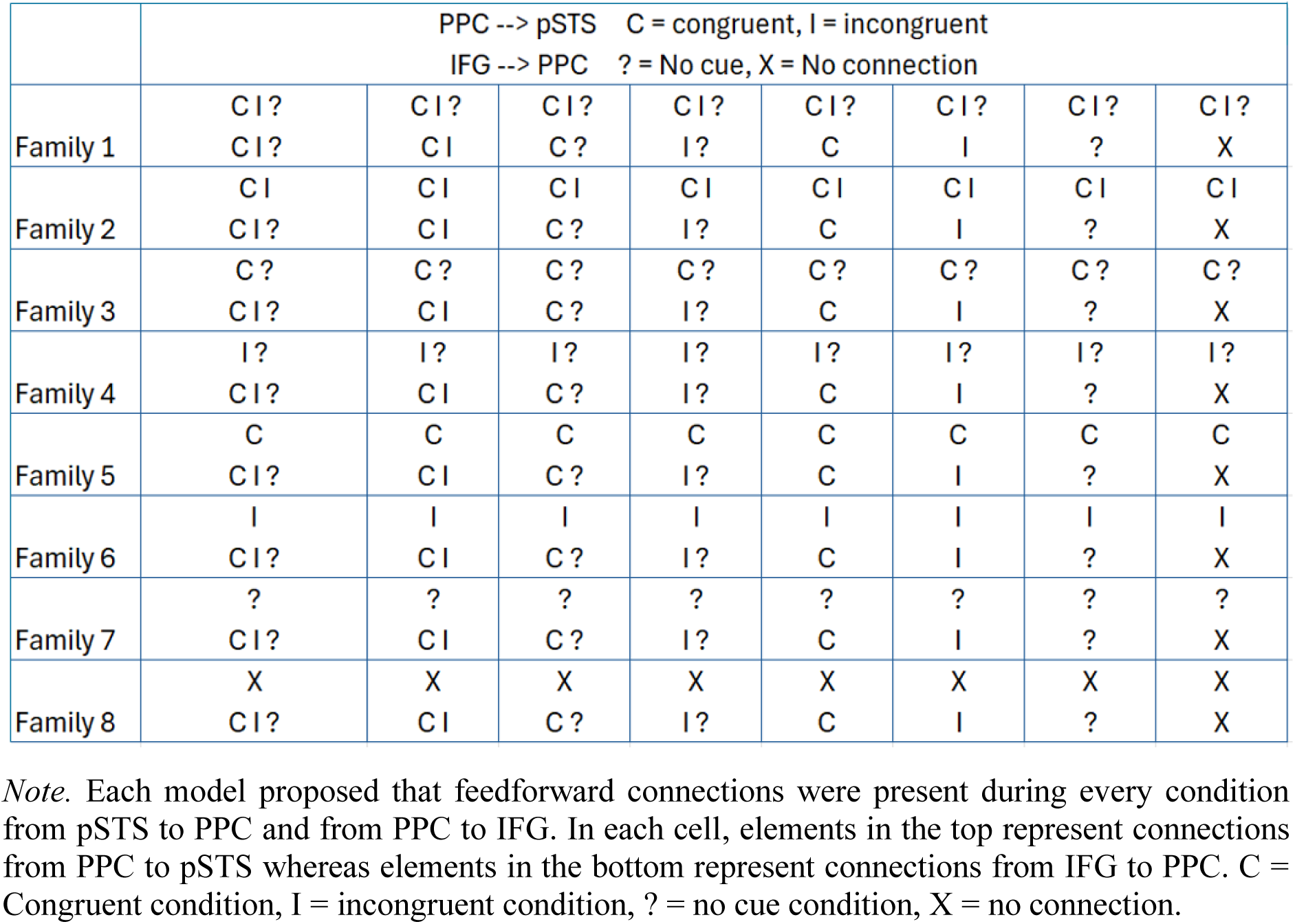
64 DCM Models.

#### 2.5.3 Multivariate Pattern Analysis (MVPA)

The data was further analyzed using MVPA, a machine-learning method. A general linear model was created in the SPM12 toolbox of MATLAB to conduct MVPA. The model included 36 regressors. 6 of these regressors were the non-stimulus periods (cue period, interstimulus interval after the cue, response period, feedback period, interstimulus interval between trials, rest period). 24 of these regressors were the trials of a run (such as kick-kick congruent trial #1, walk-kick incongruent trial #2, ?-walk no cue trial #1 etc.). The last 6 regressors were the head movements of the participant in 3 dimensions and 2 directions. The Decoding Toolbox of MATLAB was utilized for MVPA (Hebart et al., 2015). Since the experiment included m ore congruent trials than incongruent trials because of its 75% congruency nature, only 1/3 of randomly selected congruent trials were analyzed in MVPA for each participant. The chosen congruent trials spanned an equal number of kick-kick and walk-walk trials. MVPA was conducted by the searchlight method, using a sphere of 4mm radius. Cross-validation was done by the leave-one run-out method and Support Vector Machine was used as the classifier.

## 3. Results

### 3.1 Behavioral Results

The conducted behavioral analysis excluded 8 uncompleted runs across 4 participants.

#### 3.1.1 Accuracy Results

For the accuracy analysis, missed trials were treated as incorrect responses. Repeated measures ANOVA was conducted on the Greenhouse-Geisser corrected data as the assumption of sphericity was violated (p<0.05). A main effect of condition was observed (F(1.619,45.322) = 8.410, p < 0.01, η²p = 0.231) (congruent (Mean = 75.2%, SE = 2.2%), incongruent (M = 71.7%, SE = 2.9%), and no cue (M = 78.8%, SE = 1.9%)). Post hoc tests (corrected for family comparison) revealed that the accuracy of the participants was higher during the no-cue condition compared to the incongruent condition (t(28) = 4.101, p < 0.001). On the other hand, there was no significant difference neither between the congruent and no cue conditions nor between the congruent and incongruent conditions (p > 0.05 for both) (Figure 3.1).

**Figure 3.1.**
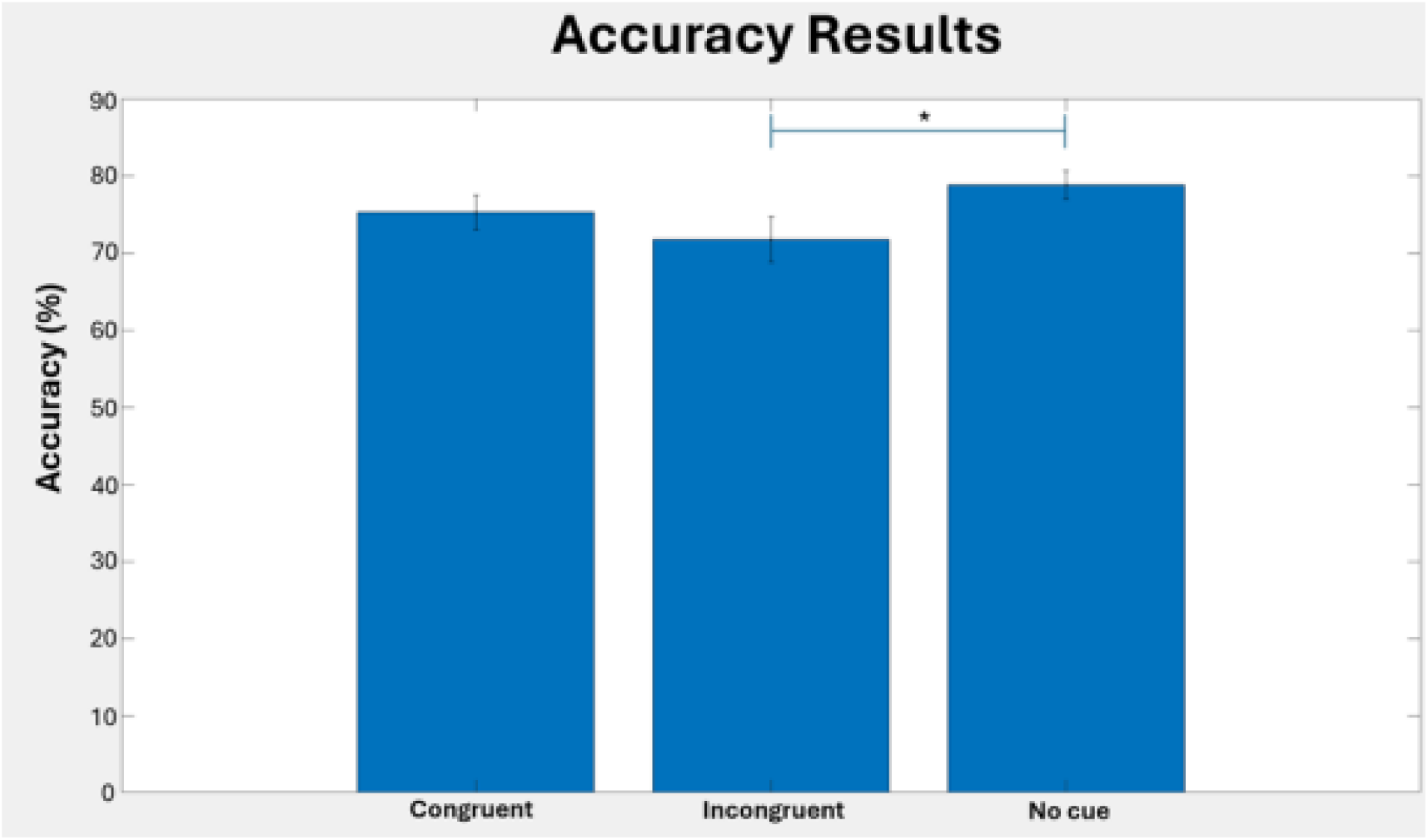
Accuracy Results.

#### 3.1.2 Reaction Time Results

Repeated measures ANOVA results showed a main effect of condition (F(2,56) = 8.284, p < 0.001, η²p = 0.228) (congruent (M = 0.572, SE = 0.023), incongruent (M=0.615, SE = 0.026) and no cue (M=0.611, SE = 0.029)). Post hoc tests (corrected for family comparison) revealed a lower reaction time during congruent condition compared to incongruent condition (t(28) = -3.677, p = 0.002) and compared to no cue condition (t(28) = -3.350, p = 0.003), whereas there was no significant difference between incongruent and no cue conditions (p > 0.05)(Figure 3.2).

**Figure 3.2.**
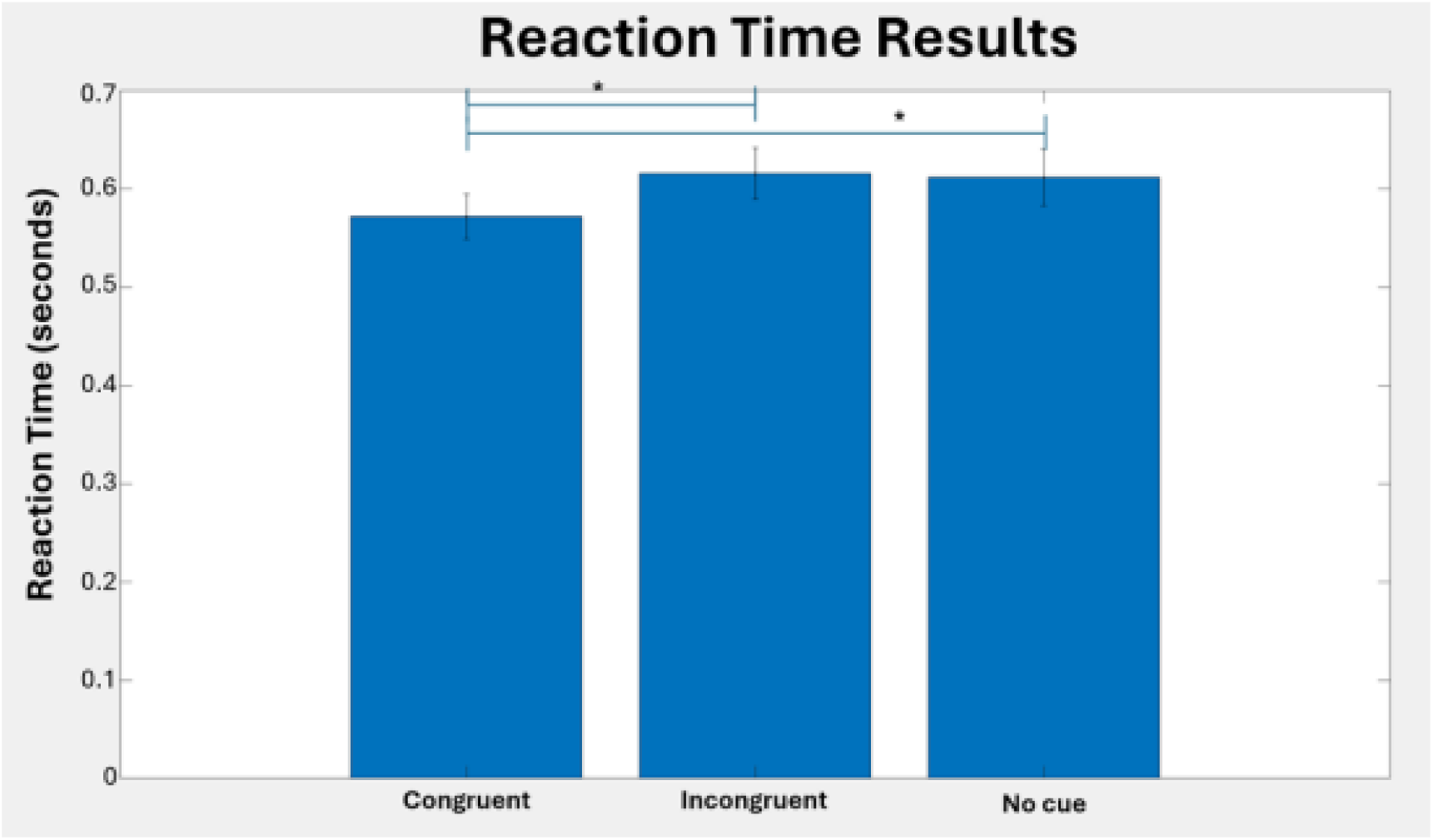
Reaction Time Results.

### 3.2 DCM Model Selection

The models were created for 8 runs of each subject. Bayesian model selection selected the first family as the winning family with an expected probability of 67.67% and an exceedance probability of 100%. Within the first family, the first model was selected as the winning model with an expected probability of 63.34% and an exceedance probability of 99.98%. The winning model stated that both types of connections (feedback and feedforward) were present between pSTS and PPC, and between PPC and IFG. Additionally, it was also suggested by the winning model that these connections were modulated by all of the conditions (congruent, incongruent, and no cue) (Figure 3.3 and Table 3.4).

**Figure 3.3.**
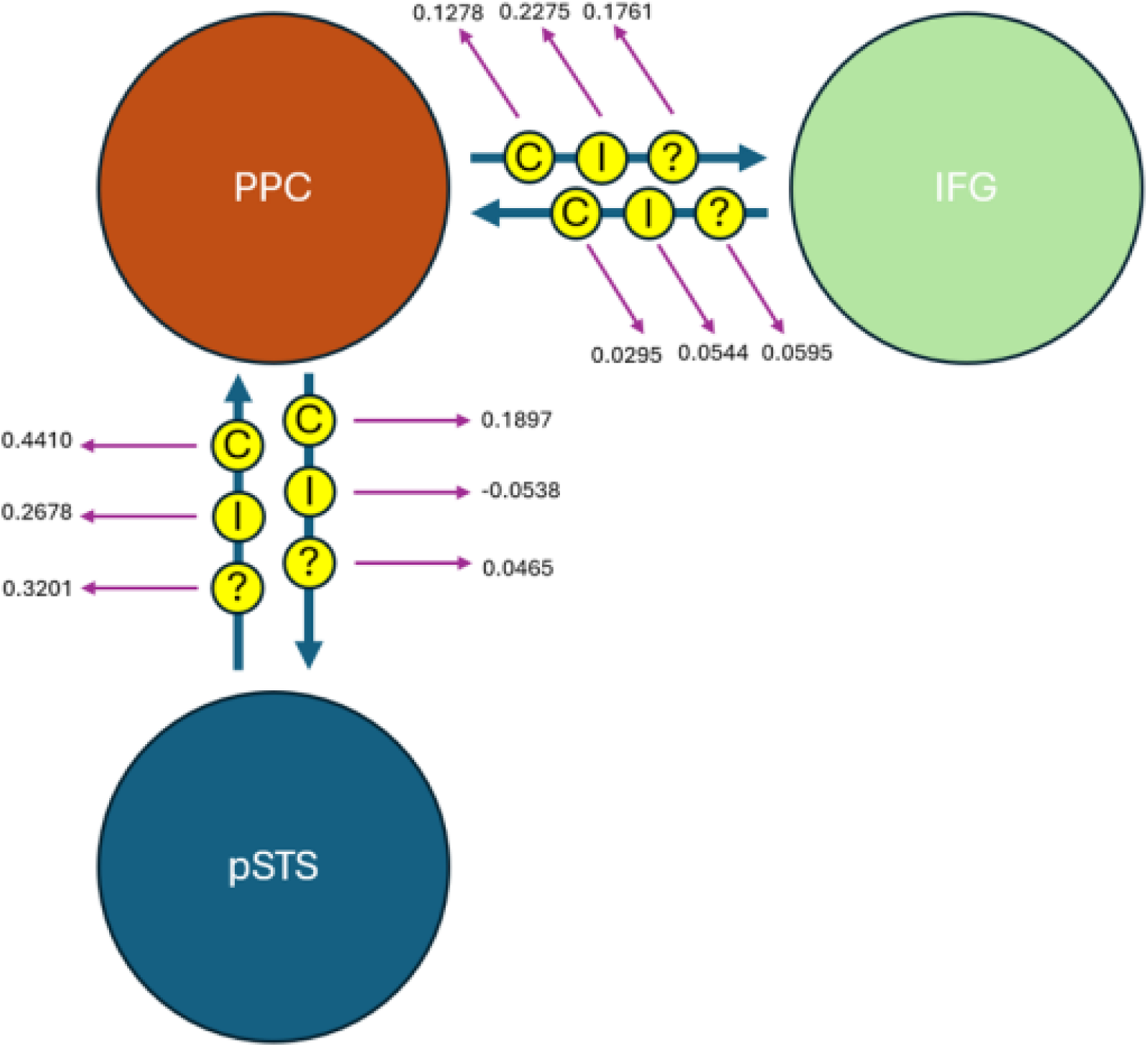
The winning model. *Note.* Modulatory connections of the model can be seen in the figure.

**Figure 3.4.**
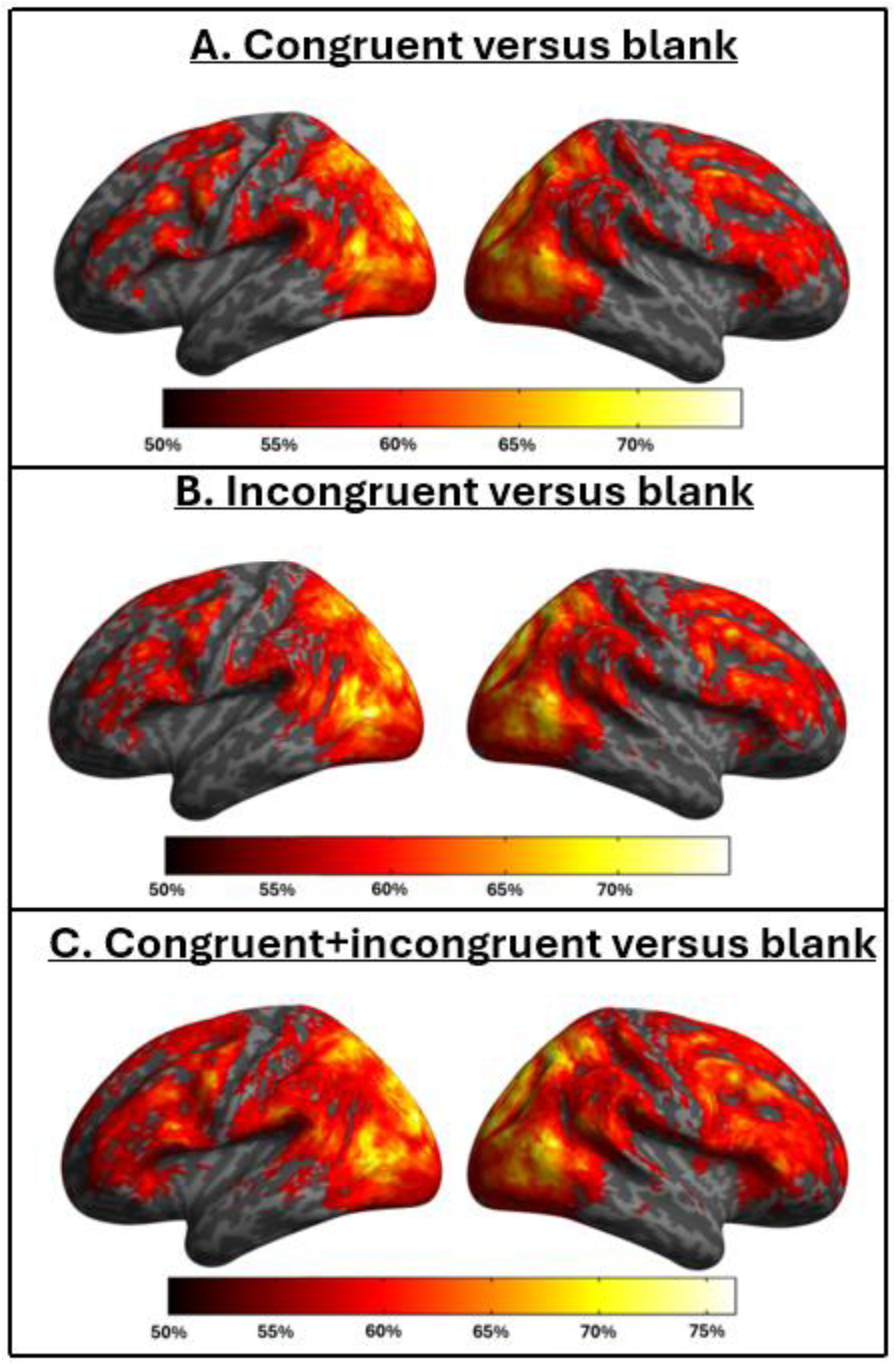
MVPA results. *Note.* **A.** Congruent and blank trials **B.** Incongruent and blank trials **C.** Congruent+incongruent and blank trials. For all figures, color scale represents classification accuracy, p < 0.05, FWE – corrected, k = 5.

**Table 3.1.**
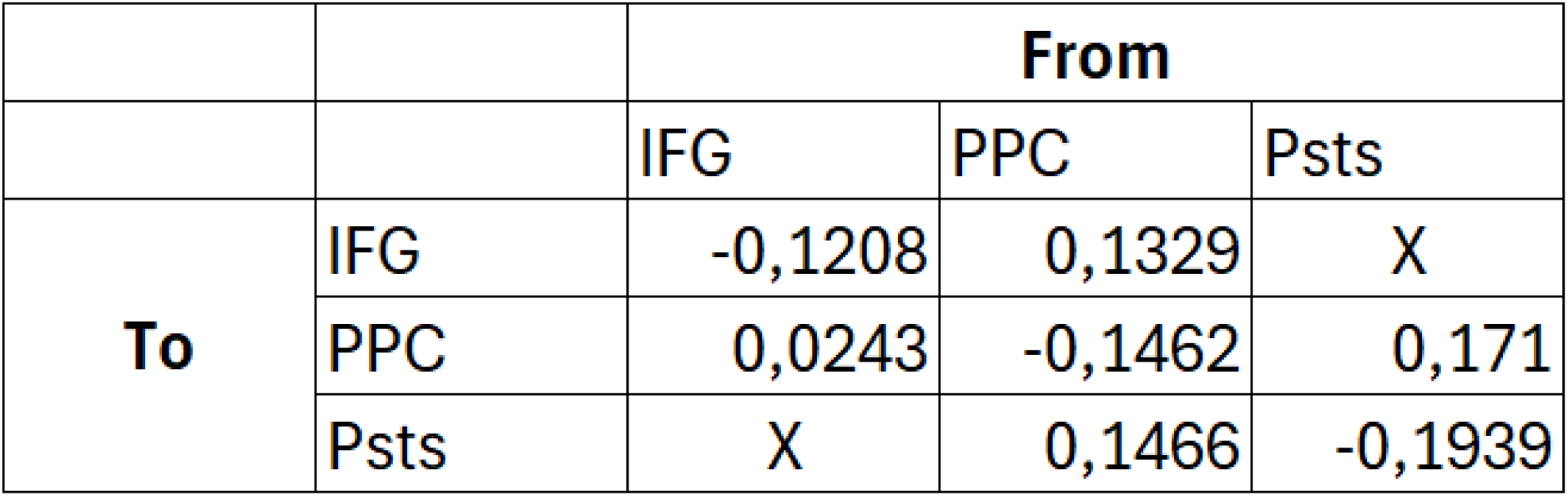
Winning Model’s Endogenous Connections.

### 3.3 MVPA

5 participants were excluded from MVPA for the following reasons: 2 participants’ data were not usable because of head motion and 3 participants’ data were not usable because of a technical problem with the scanner. 7 condition pairs were created for this analysis: Congruent and incongruent, congruent and no cue, incongruent and no cue, blank and congruent, blank and incongruent, combination of congruent and incongruent trials (referred as congruent+incongruent) and blank, congruent+incongruent and no cue. The pairs that did not contain the blank condition failed to show any significant result (p > 0.05, FWE-corrected, k = 5 for all). On the other hand, blank and congruent (Figure 3.3), blank and incongruent (Figure 3.4), and blank and congruent+incongruent (Figure 3.5) condition pairs showed significant results (p < 0.05, FWE-corrected, k = 5 for all)(Table 3.3).

**Table 3.2.** MVPA Local Maxima Values.

|  | MVPA Results |  |  |  |  |  |
| --- | --- | --- | --- | --- | --- | --- |
|  | Left |  |  | Right |  |  |
|  | Congruent | Incongruent | Congruent+<br>Incongruent | Congruent | Incongruent | Congruent+<br>Incongruent |
| PPC |  |  |  | x=24<br>y=-72<br>z=56 | x=20 y=-76<br>z=54 | x=20 y=-76<br>z=54 |
| MSFG | x=-6 y=50<br>z=6 |  |  | x=2 y=68<br>z=6 |  |  |
| Caudate | x=-10<br>y=18 z=-4 |  |  |  |  |  |
| Thalamus<br>Proper |  | x=-4 y=-0<br>z=-8 | x=-6 y=-24<br>z=14 |  | x=12 y=6<br>z=6 |  |
| MFG |  | x=-34 y=54<br>z=-4 |  |  |  |  |
| PCgG |  |  |  |  | x=2 y=-42<br>z=4 |  |
| FuG |  |  | x=-30 y=-40<br>z=-16 |  |  |  |

## 4. Discussion

The current study aimed to investigate the influence of expectation on biological motion perception at the cortical level. The participants were given a decision-making task during which they were given cues that created correct or wrong predictions about the action that they were about to see while fMRI activity was recorded. The data was analyzed using dynamic causal modeling technique to understand the effective connectivity within nodes of the action observation network (pSTS, PPC, IFG). A multivariate pattern analysis was also conducted to investigate whether nodes of AON can successfully decode between expected and unexpected actions.

The behavioral accuracy results show that the participants tend to be more accurate under the lack of a cue compared to the presence of a false cue. The participants performed better during the no cue condition compared to the incongruent cue condition, whereas the congruent condition did not differ from other conditions. Thus, the results show that false prior information about the stimulus impairs the performance of the participants. This effect successfully demonstrates the effect of top-down processes on biological motion perception. In line with the accuracy results, reaction time data also demonstrates the effect of top-down processes on biological motion perception such that the participants’ perception was the fastest when a correct cue was presented to them compared to other conditions. Ultimately, the behavioral findings suggest an enhanced processing of biological motion when the cue is congruent. Therefore, behavioral results support the previous studies in the perception literature that show a hindering effect of false cues and a facilitating effect of congruent cues on perception (Kok et al., 2014; Kok et al., 2012; Kok et al., 2013; Urgen & Boyaci, 2021; Elmas et al., 2024; Loof et al., 2016; Aitken et al., 2020; Chen et al., 2019; Margoni et al., 2024).

In terms of brain imaging, the DCM results show significant effective connectivity within nodes of AON. In light of the current literature, 64 models were created. The winning model, selected by the Bayesian Model Selection procedure, argues that the information enters pSTS and follows a feedforward path from pSTS to PPC, and from PPC to IFG. Additionally, the model acknowledges the presence of feedback connections during all three congruence conditions from IFG to PPC, and from PPC to pSTS. The winning model provides three essential contributions to the biological motion and prediction literature. Firstly, the model successfully demonstrates that AON consists of both feedback and feedforward connections. This finding supports Urgen & Saygin’s (2020) mismatch model and also the anatomical studies that show PPC having reciprocal connections with the other two nodes of the network (Luppino et al., 1999; Seltzer & Pandya, 1994; Rushworth et al., 2006; Igelstrom & Graziano, 2017). More importantly, by showing the presence of feedback connections, the model successfully challenges bottom-up models of biological motion perception. It is worth noting that the winning model does not discard the importance of bottom-up connections, but proposes that biological motion perception requires both bottom-up and top-down information. Secondly, the model states that AON activity depends on the generation and violation of predictions. Although the role of attention as a top-down factor in biological motion perception has been established, the current study provides empirical evidence for the role of expectations, while also supporting the view of AON as a predictive coding system (Friston, 2010). Thirdly, the winning model becomes one of the few studies that propose the presence of inhibitory connections within AON. More specifically, it successfully shows PPC has an inhibitory feedback effect on pSTS when expectations are violated. Since expectation violation with simple stimuli (such as Gabor patches) causes a similar inhibitory feedback signal as well, this successfully generalizes the findings of prediction literature to biological motion, a highly complex and social stimulus (Kok et al., 2014; Kok et al., 2012, Kok et al., 2013). Ultimately, the study proposes a prior information model for biological motion perception literature, building upon the previous biological motion and predictive coding studies.

In addition to the DCM results, the FWE-corrected searchlight MVPA results showed a significant and strong decoding for biological motion in the occipital cortex, pSTS, PPC, and IFG. All of these regions are successfully able to decode between stimulus and non-stimulus conditions, demonstrating their essential role in biological motion and action perception. Moreover, the study also managed to provide strong evidence of fusiform gyrus’ biological motion selectivity by demonstrating successful decoding in the region (Vaina et al., 2001). Nevertheless, the MVPA results did not align with the behavioral findings of this study or previous literature, as they failed to demonstrate successful decoding among stimulus conditions (Kok et al., 2012; Alink et al., 2010).

It is possible that the lack of a significant MVPA result in stimulus conditions was caused by the primary limitation of the study, and that discussing this limitation is essential to better interpret the results. Although 75% congruence rate was decided to be optimal for the study, this rate has caused it to suffer from a low number of trials for certain conditions. In order to achieve 75% congruence, the incongruent condition was designed with three times fewer trials than the congruent condition. Consequently, no cue and blank conditions also had a low trial count. As the same number of trials across conditions were required for MVPA analysis, ⅓ of congruent trials had to be randomly selected, forcing the analysis to be conducted with a relatively low number of trials. Thus, it is highly possible that the lack of a significant result might have been caused by the low number of trials. Although increasing the number of trials was possible, this option was not pursued to maintain a reasonable experiment duration and to prevent potential learning and fatigue effects.

Another MVPA finding worth mentioning is the local maxima observed in the prefrontal cortex and striatum. During the study, the participants were given a decision-making task. Hence, activation in the prefrontal cortex was expected as the region plays a crucial role in decision-making and biological motion perception (Jastorff & Orban, 2009; Peuskens et al., 2005; Lesourd et al., 2022; Orban, 2018; Grezes & Decety, 2001; Saygin et al., 2004; Miller & Cohen, 2001; Bechara et al., 1997; Sanfey et al., 2003). However, the observed activation was stronger than anticipated. This result suggests the possibility that prediction errors may have been generated during blank trials, given that a crucial role of the prefrontal cortex is the generation of prediction errors (Asaad & Eskandar, 2011; Miller & Cohen, 2001). This idea is further supported by the local maxima observed in the striatum, another region crucial for the generation of prediction errors (Schultz, 1998; Schiffer et al., 2012; Schiffer & Schubotz, 2011). It is believed that the task created a second, unintended expectation: Since 83.33% of the trials included a biological motion, the participants expected to see a biological motion in every trial, which resulted in a violation of this expectation during blank trials. This result successfully supports prediction literature by showing the role of the prefrontal cortex and striatum on the generation of prediction errors and contributes to the biological motion literature by showing the presence of top-down mechanisms in biological motion perception. However, it is outside of the scope of the present study as predicting the presence of the stimulus is a task-relevant prediction, whereas the study aims to investigate task-irrelevant predictions.

Additionally, the current study also opens up numerous possibilities for future experiments. The study has succeeded in building a biological motion perception model with both feedback and feedforward connections. As this was the first study to do so, 2D point-light displays of non-communicative actions were used to have a highly controlled stimulus. Future studies can build on the findings of the current study by using more varied stimuli. Biological motion literature has demonstrated that communicative actions (the actions that are aimed at another agent) result in higher PPC activation. Similarly, it was shown by multiple studies that real-life videos of actions cause a stronger activation in the action observation network as they better reflect daily life scenarios (Beauchamp et al., 2003; Jastorff et al., 2016; Hars et al., 2011). Based on the result of this study, it is expected that using stimuli that cause stronger AON activation is likely to show significant MVPA results as well. Lastly, the current study can be replicated with different congruence rates or different cues to further generalize the findings of current and any future study.

In conclusion, the study at hand contributed to the biological motion literature by building a model that explains the role of prior information and top-down processes on biological motion perception, while also supporting the existing literature by showing the role of action observation network and fusiform gyrus in biological motion perception. Additionally, the study also contributed to prediction literature by showing the brain regions that are involved in prediction paradigms with socially meaningful stimuli.

## Acknowledgments

The study was supported by a TUBITAK 3501 grant awarded to Burcu A. Urgen (Grant No: 119K654). The authors would like to thank Hilal Nizamoğlu, Aysu Koç, Aslı Eroğlu, Zelal Eltaş and Yunus Emre Türkmen for their help with data collection.

